# The Past Sure Is Tense: On Interpreting Phylogenetic Divergence Time Estimates

**DOI:** 10.1101/113720

**Authors:** Joseph W. Brown, Stephen A. Smith

## Abstract

Divergence time estimation — the calibration of a phylogeny to geological time — is an integral first step in modelling the tempo of biological evolution (traits and lineages). However, despite increasingly sophisticated methods to infer divergence times from molecular genetic sequences, the estimated age of many nodes across the tree of life contrast significantly and consistently with timeframes conveyed by the fossil record. This is perhaps best exemplified by crown angiosperms, where molecular clock (Triassic) estimates predate the oldest (Early Cretaceous) undisputed angiosperm fossils by tens of millions of years or more. While the incompleteness of the fossil record is a common concern, issues of data limitation and model inadequacy are viable (if underexplored) alternative explanations. In this vein, Beaulieu *et al*. (2015) convincingly demonstrated how methods of divergence time inference can be misled by both (i) extreme state-dependent molecular substitution rate heterogeneity and (ii) biased sampling of representative major lineages. These results demonstrate the impact of (potentially common) model violations. Here, we suggest another potential challenge: that the configuration of the statistical inference problem (i.e., the parameters, their relationships, and associated priors) alone may preclude the reconstruction of the paleontological timeframe for the crown age of angiosperms. We demonstrate, through sampling from the joint prior (formed by combining the tree (diversification) prior with the calibration densities specified for fossil-calibrated nodes) that with no data present at all, that, an Early Cretaceous crown angiosperms is rejected (i.e., has essentially zero probability). More worrisome, however, is that, for the 24 nodes calibrated by fossils, almost all have indistinguishable marginal prior and posterior age distributions when employing routine lognormal fossil calibration priors. These results indicate that there is inadequate information in the data to overrule the joint prior. Given that these calibrated nodes are strategically placed in disparate regions of the tree, they act to anchor the tree scaffold, and so the posterior inference for the tree as a whole is largely determined by the pseudo-data present in the (often arbitrary) calibration densities. We recommend, as for any Bayesian analysis, that marginal prior and posterior distributions be carefully compared to determine whether signal is coming from the data or prior belief, especially for parameters of direct interest. This recommendation is not novel. However, given how rarely such checks are carried out in evolutionary biology, it bears repeating. Our results demonstrate the fundamental importance of prior/posterior comparisons in any Bayesian analysis, and we hope that they further encourage both researchers and journals to consistently adopt, this crucial step as standard practice. Finally, we note that the results presented here do not refute the biological modelling concerns identified by Beaulieu *et al.* (2015). Both sets of issues remain apposite to the goals of accurate divergence time estimation, and only by considering them in tandem can we move forward more confidently. [marginal priors; information content; diptych; divergence time estimation; fossil record; BEAST; angiosperms.]

---

“Molecular clocks are not up to the job, but neither is the fossil record.” Donoghue and Benton (2007)

Divergence time estimation from molecular genetic sequences is fraught with uncertainty. The errors involved in routine phylogenetic reconstruction (suboptimal alignments, inadequate substitution models, insufficient taxon/gene sampling, real gene tree/species tree conflict, etc.) are compounded by assumptions required to transform a phylogram (in units of expected number of substitutions per site) into a chronogram (in units of geological time): (i) an appropriate model of substitution rate heterogeneity among lineages and across time, and (ii) temporal calibrations, generated from the fossil (or biogeographic) record, used to inform and constrain the extent of rate variation.

It is therefore not surprising that there are discrepancies between inferred molecular genetic and paleontological timescales. However, while many disagreements are minor and may innocuously be attributed to insufficient sampling (genes, taxa, or fossils), others are so severe and consistent that they cast serious doubt on the appropriateness of molecular clock models, the fossil record, or both. One prominent example concerns placental mammals, where molecular estimates (e.g., Meredith *et al.* 2011) of the crown age almost double those from the fossil record (e.g., O’Leary *et al.* 2013), obscuring the role of the K-Pg mass extinction on the evolutionary trajectory of this group. Another conspicuous example, also spanning the K-Pg boundary, is crown birds (Neornithes; Ksepka *et al.* 2014), where (re)analyses have repeatedly led to incongruous inferred evolutionary timeframes (e.g., Ericson *et al.* 2006 vs. Brown *et al.* 2007; Jarvis *et al.* 2014 vs. Mitchell *et al.* 2015; Prum *et al.* 2015).

However, perhaps the best exemplified recalcitrant node in terms of absolute age is that of crown angiosperms (flowering plants), where molecular clocks pervasively infer a Triassic age (e.g., Bell *et al.* 2010; Smith *et al.* 2010; Zeng *et al.* 2014; Beaulieu *et al.* 2015; Foster *et al.* 2017; (Sauquet *et al.*, 2017); see a comprehensive review of estimates in Magallón *et al.* 2015) while the oldest undisputed fossil remains are restricted to the Early Cretaceous (136 Ma; Brenner 1996; see a recent review in Herendeen *et al.* 2017). Moreover, Magallón *et al.* (2015), applying a model of uniform random fossilization (Marshall 2008) to 136 fossils, inferred an upper bound on the origin of crown angiosperms of just ~140 Ma Finally, a Bayesian modelling of fossil preservation estimated that crown angiosperms originated 151.8-133.0 Ma (Silvestro *et al.* 2015). The age of this one node, more than any other, has seriously called into question the utility of both molecular clock models and the fossil record.

Numerous reasons have been put forth to explain the disparity of molecular and paleontological timescales. On the one hand there are valid concerns with the fossil record. By their nature, fossils *must* postdate the origin of taxa, meaning that molecular estimates *should* predate those from the fossil record. Furthermore, it is clear that the fossil record is non-uniform in both space and time (Holland 2016), such that for some taxa it may prove impossible to ever have a tight correlation with molecular clock estimates. However, ‘absence of evidence’ vs. ‘evidence of absence’ is a complex matter, so a more productive avenue to pursue may be that of considering molecular approaches. While it has been speculated that molecular clocks might ‘run fast’ during radiations (thereby misleading clocks into inferring a long period of time has occurred; Benton 1999), this has no empirical support. However, it is known that substitution model mis-specification can mislead divergence time estimation (Phillips 2009; Schenk and Hufford 2010) and ultimately downstream analyses (Revell *et al.*, 2005). Likewise, mis-specification of relaxed clock models may also lead to inaccurate results (Dornburg *et al.* 2012; Worobey *et al.* 2014; Duchene and Ho 2014). Being only semi-identifiable, molecular clock methods require, in addition to molecular sequence data, calibration from the fossil record, so appropriate calibration use is critically important (Inoue *et al.* 2010; Sauquet *et al.* 2012; Warnock *et al.* 2012; Magallón *et al.* 2013; Duchene *et al.* 2014; Zhu *et al.* 2015; Barba-Montoya *et al.* 2017; Warnock *et al.* 2017). Finally, Beaulieu *et al.* (2015) recently demonstrated through simulation two further ways where molecular clocks might be misled: (i) through extreme state-dependent molecular substitution rate heterogeneity, and (ii) biased sampling of representative major lineages. Both of these issues are primarily instances of model violation. While these are all valid concerns, we explore below a further non-biological possibility that may unknowingly be at play in many data analyses.

## Diptych: A Metaphor For Data Analysis?

A diptych is a device commonly used in western art and literature. It consists of paired, complementary works, in the artistic tradition typically two images joined at a hinge (e.g., Fig. 1). The function of a diptych is to reciprocally illuminate the component images, ideally revealing some more holistic concept or narrative. It is this feature that suggests an association with Bayesian data analysis. It is *de rigueur* in any Bayesian analysis to carefully compare paired prior and posterior parameter distributions to gauge how information content (via the likelihood) drives the results, as well as to establish the sensitivity of inferences to prior specifications. A diptych is, we argue, therefore a useful metaphor for describing the process of changes in belief in parameter values from the prior (*before* data have been observed) to the posterior (*after* data have been observed). We note that the phylogenetic systematics community has been generally lax in this respect, despite available Bayesian software packages making such reflections straightforward. While the recommendation of comparing prior and posterior parameter distributions is by no means novel (rather, it is a general statistical concern), the rarity with which it is carried out in evolutionary biology bears emphasis. In this vein, we hope the diptych metaphor will prove useful in generating discussions. We argue that this is especially important in divergence time estimation analyses, as it is generally unappreciated by empirical practitioners that for fossil-calibrated nodes there are *three* sets of distributions to consider. In addition to the temporal fossil calibration prior specified by the investigator (the ‘user prior’) and the resulting marginal posterior distribution, there exists an intermediate distribution, the marginal prior (also called the ‘effective’ or ‘joint’ prior by some authors), which is formed by the interaction among user priors and the underlying ‘tree prior’ (for nodes not directly calibrated by a fossil; typically a birth-death prior). Here we turn our attention to these distributions to see what, if anything, we can glean about the age of crown angiosperms.

**FIGURE 1.**
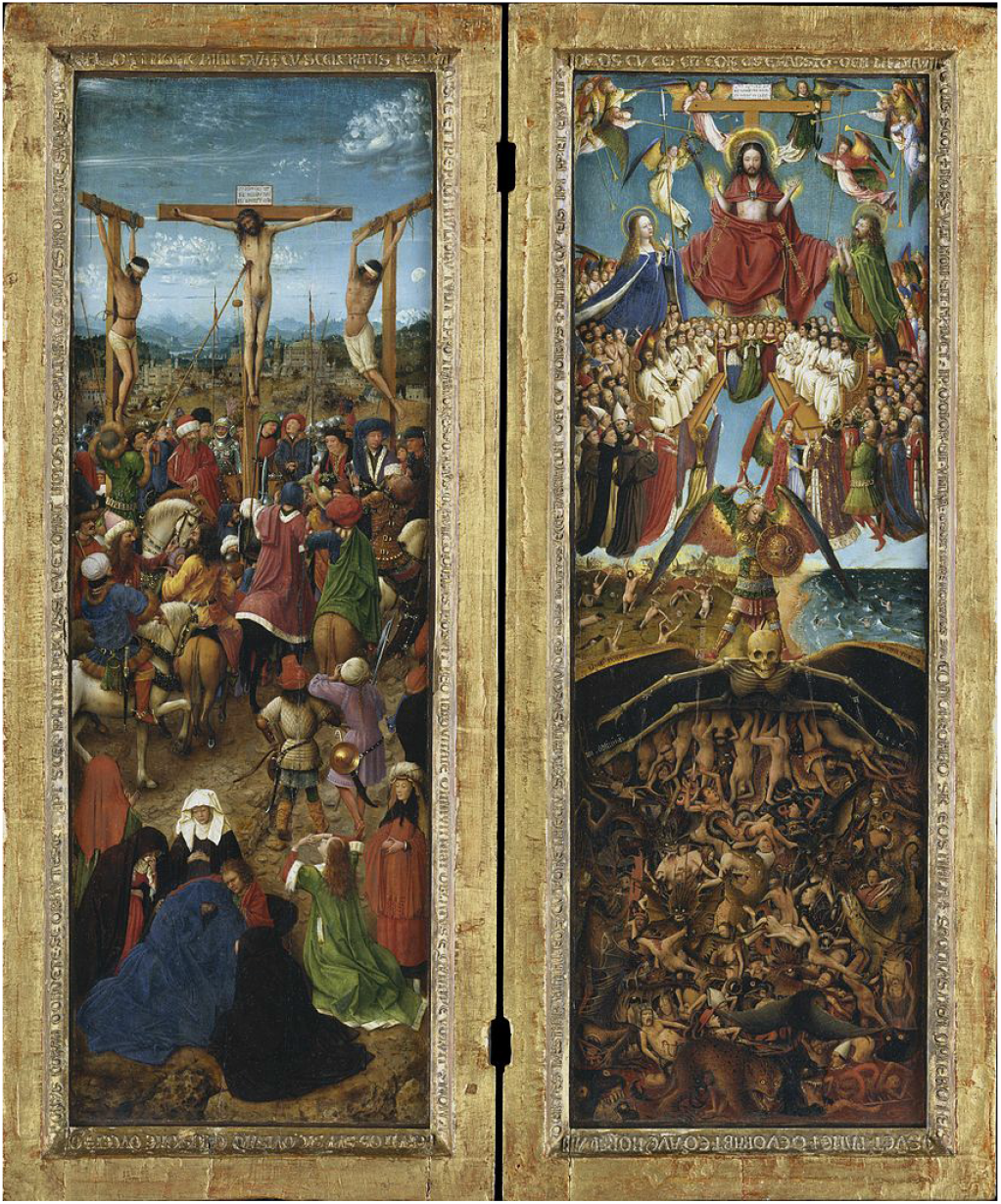
Crucifixion and Last Judgement diptych by Jan van Eyck c. 1430—1440 (Public domain). The individual images, depicting scenes from the Christian Bible, can be viewed in isolation, but the work as a whole conveys a coherent narrative only when considering the component pieces together. The temporal aspect of this particular work, with contrasting before (left) and after (right) titular events, mirrors the temporal process in Bayesian data analysis where belief in parameter values is updated from prior to posterior distributions.

## A Re-Reanalysis Of The Age Of Angiosperms

We reanalyzed the data set provided by Beaulieu *et al.* (2015). The molecular alignment consists of four genes (chloroplast: *atp*B, psbB, and *rbcL*; nuclear: 18S) for 124 taxa including 91 angiosperms representing all extant orders (data file provided in the Supplementary Material; see also data from the original paper available on Dryad at http://dx.doi.org/10.5061/dryad.629sc). Sampling was originally designed specifically for dating the origin of angiosperms by allowing the placement of 24 (15 angiosperm) fossil calibrations from across landplants. All dating analyses reported here, like those in the original paper, were performed using the uncorrelated lognormal (UCLN) clock model and birth-death tree prior in BEAST v1.8.2 (Drummond *et al.* 2006; Drummond and Rambaut 2007) using the CPU implementation of the BEAGLE v2.1.2 library (Ayres *et al.* 2012).

We regenerated the posterior results of Beaulieu *et al.* (2015) by employing the original lognormal user priors specified in Table 1 and the following analysis settings: 3 replicate analyses of 50 million generations, sampling every 1000 generations. As in the original Beaulieu *et al.* (2015) paper we fixed the tree topology (their Figure 1). These analyses were re-run without any data (i.e., sampling from the marginal prior). Because these latter analyses were not as computationally demanding, we attempted to precisely estimate the breadth of marginal prior distributions (through minimizing Monte Carlo error) by running analyses in duplicate for 1 billion generations each, sampling every 10 thousand generations. Finally, we replicated all analyses but replaced the lognormal user prior calibrations from Table 1 with ‘extreme exponential’ distributions with a mean of 1.0 (offset by fossil ages); such distributions lend the utmost credence to the fossil record, as 95% of the prior masses lie within 3 Ma of the relevant fossil ages. Importantly, all analyses (posterior and prior-only) were initialized with chronograms wherein crown angiosperms originated ~140 Ma (i.e., consistent with the prescription of Magallón *et al.* 2015). Analyses were initialized in this way to ensure that the MCMC sampler had a definite chance of sampling younger ages (something that is not guaranteed for finite chain lengths).

**TABLE 1.**
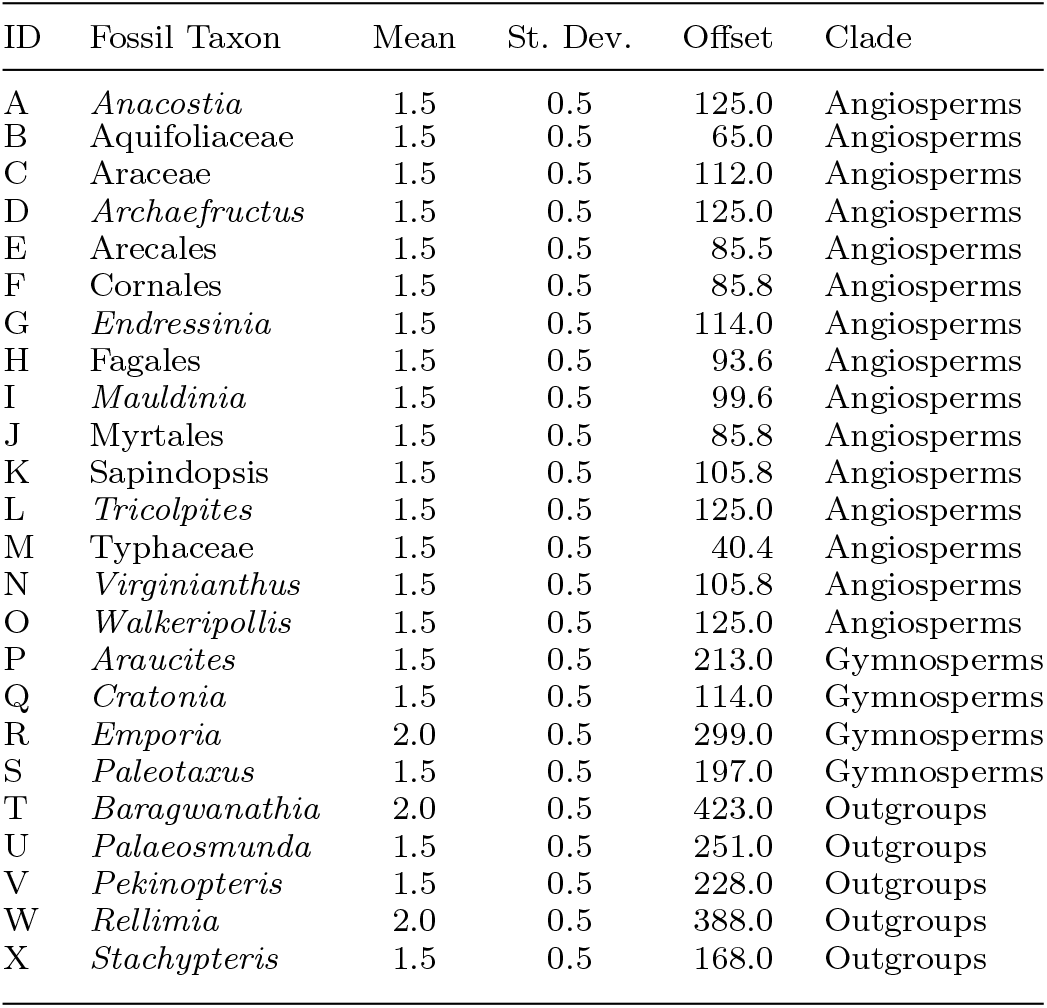
Lognormal fossil calibration parameters as originally defined in Beaulieu *et al.* 2015.

Notes: ‘Offset’ denotes the age of the oldest undisputed fossil in Ma Mean and standard deviation are given in log space. Distributions with a mean of 1.5 have 95% of the prior mass within 10.2 Ma of the fossil age, while those with a mean of 2.0 have 95% of the prior mass within 16.82 Ma of the fossil age. As we focus on the crown angiosperm node, we note only whether the fossils fall within the clade, within the sister clade, or outside. Interested readers can see Beaulieu *et al.* (2015) for a justification of the placement and forms of these calibrations.

We focus here on the data (taxon, gene, and fossil sampling) and settings (model and priors) of Beaulieu *et al.* (2015) because they are representative of a standard dating analysis. However, to demonstrate that our own results are not restricted to this particular data set, we also reanalyze the data set of Magallón *et al.* (2015), albeit to a more limited degree because of computational requirements. This data set consists of 5 genes (chloroplast: *atp*B, rbcL and *matK;* nuclear: 18S and 26S) for 799 taxa (792 angiosperms) and 121 fossil calibrations. We note that the original analysis by Magallón *et al.* (2015) included 137 fossil-based calibrations. Our inclusion of 121 fossil calibrations is the result of the file, shared by the original authors, having 16 fewer calibrations. As we are more interested in contrasting prior and posterior distributions, and demonstrating parallel patterns across data sets, these results are still valuable. All analyses employed the UCLN model as above and a fixed tree topology (their Figure 3). The posterior results of Magallón *et al.* (2015) were regenerated by running 3 replicate analyses of 100 million generations, sampling every 5000 generations. These analyses included a uniform calibration prior (139.5-136 Ma) on the age of crown angiosperms applied by the original authors. To assess the influence of this single prior, 4 replicate analyses of 50 million generations were performed without the prior. Finally, as above, analyses sampling from the marginal prior (i.e., without any data) were performed for both sets of fossil calibrations (i.e., with and without the crown angiosperm temporal constraint), each with 4 replicates of 50 million generations. Due to computational restrictions, we did not explore the use of the ‘extreme exponential’ calibration priors for this data set, and instead restrict analyses to using the original fossil calibration priors from Magallón *et al.* (2015). For a thorough exploration of calibration prior sensitivity, see Foster *et al.* (2017).

Prior to summarization, we identified a 10% sample burnin as being conservative (i.e., convergence was achieved prior to this in all analyses). MCMC log files from each set of replicated analyses were concatenated while removing this 10% sample burnin using the pxlog program from the phyx package (Brown *et al.* 2017). All results were processed in R v3.3.2 (R Core Team 2016) and were visualized using ggplot2 v2.2.1 (Wickham 2009) and code adapted from phyloch v1.5-3 (Heibl 2008).

## The Inaccessibility Of An Early Cretaceous Crown Angiosperms

As in the original Beaulieu *et al.* (2015) paper we were unable to recover a posterior age estimate for crown angiosperms that corresponded with the prevailing paleontological timeframe, even when employing overly precise exponential fossil calibration user priors. However, when considering the diptych interpretation by examining the joint marginal prior, it is clear that we need not invoke modelling complications (e.g., due to structured excessive rate heterogeneity) to explain the results. Rather, when running the analysis without any data (Fig. 2), we see that an Early Cretaceous crown angiosperms is precluded based on the configuration of the statistical problem alone (i.e., the set of parameters, their relationships, and the form and interaction of their associated priors). From the trace plots (Fig. S1) we see that the parameter regarding the age of crown angiosperms departs immediately from ~140 Ma to >200 Ma In no instance did the MCMC samplers ever return to a ‘young’ age of angiosperms. The youngest post-burnin ages for the prior and posterior analyses for the original lognormal calibration priors were 185.9 Ma and 192.0 Ma, respectively (181.1 Ma and 176.0 Ma for the exponential calibration priors). We note that these findings do not have to do with any peculiarity of the Beaulieu *et al.* (2015) data set as the results generated from the Magallón *et al.* (2015) data set without the uniform prior on crown angiosperms confirms the findings (minimum post-burnin ages for the prior and posterior analyses are 226.1 Ma and 197.2 Ma, respectively). In fact, the Magallón *et al.* (2015) data set, containing far more data (genes, taxa, and fossils) generated the oldest mean posterior estimate (249.7 Ma vs. 233.0 Ma for the original Beaulieu *et al.* (2015) priors vs. 213.3 Ma for the same data set using exponential calibration priors). Sauquet *et al.* (2017) found similar results when removing this fossil constraint.

**FIGURE 2.**
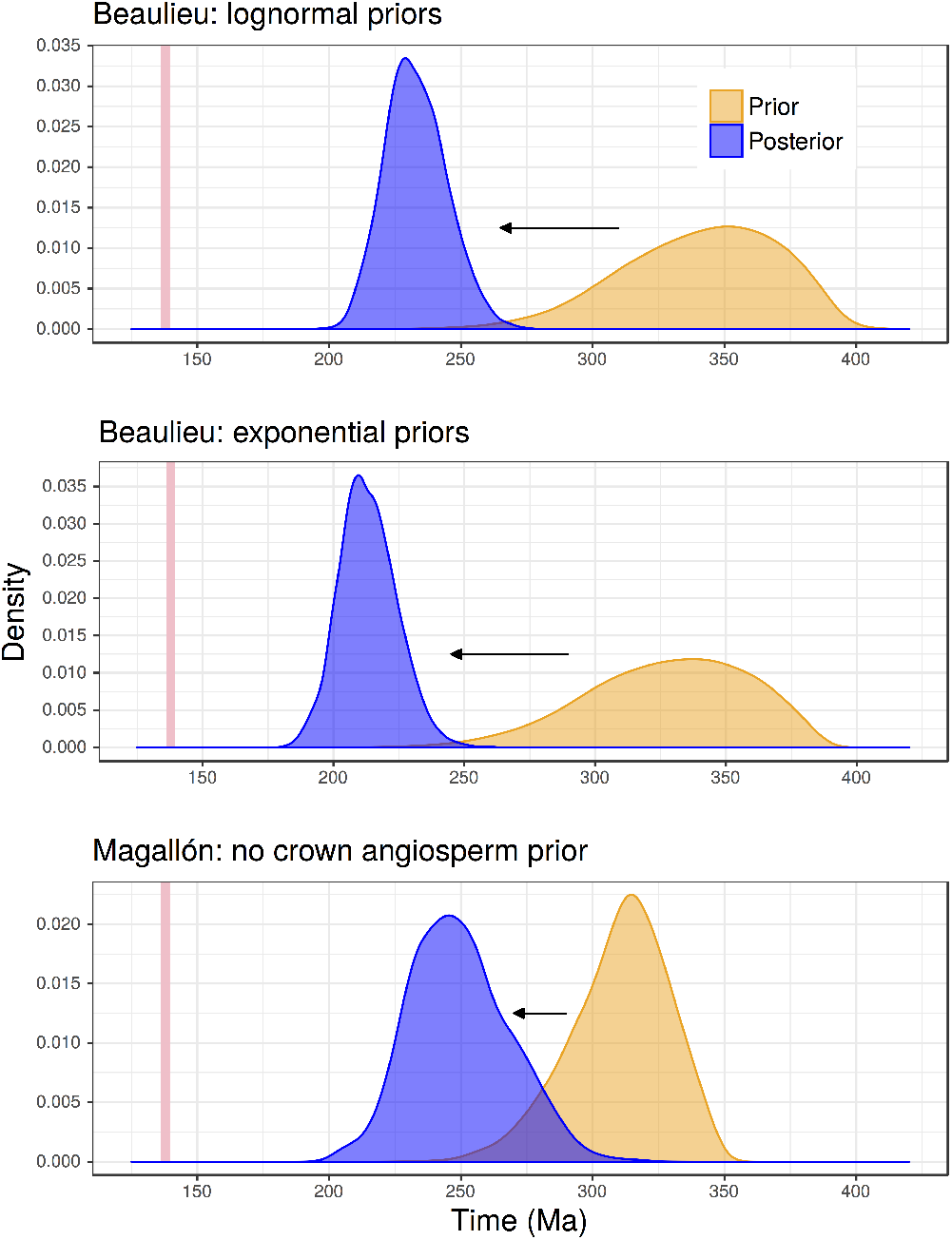
Diptychs comparing marginal prior (right) and posterior (left) distributions for the age of crown angiosperms. The top panel displays results using the original lognormal fossil calibration priors with the Beaulieu *et al.* (2015) data set, while the middle panel uses the exponential priors for the same data set. The bottom panel displays results for the Magallón *et al.* (2015) data set. Note that none of the analyses include a user prior for this node. For reference the uniform prior (139.5-136 Ma; vertical bar) used by Magallón *et al.* (2015), reflecting the paleontological estimate, is shown.

While rejecting an Early Cretaceous origin based on the marginal prior alone, both data sets *do* seem to contain signal relevant to the age of crown angiosperms, as the marginal posterior estimates are shifted significantly younger than the marginal prior (Fig. 2). This raises the question: if true, what kind/amount of data would be required to recover an Early Cretaceous age for crown angiosperms? Ultimately this comes down to quantifying phylogenetic ‘information content’. Intriguing possibilities to addressing this sort of question therefore lie in Shannon information theory (Shannon 1948), a field which the systematics community has largely ignored. Recently Lewis *et al.* (2016) made substantial strides forward by applying this theory (in the discrete case) towards assessing the information content of prior and posterior tree topology distributions. Applications to the continuous case, however, are much more difficult, and no theory (let alone software) currently exists to address the present problem.

## Priors And Posteriors: A Diptych In Three Parts

Above we introduced ‘diptych’ as a potentially useful metaphor for interpreting Bayesian analytical results. The paired nature of a diptych mirrors the before (prior) and after (posterior) reflection on what has been learned about probable parameter values.

The metaphor is slightly more complicated for some parameters involved in divergence time analyses. Nodes not explicitly calibrated by fossil data (henceforth, ‘uncalibrated’ nodes) still require an age prior; this is provided by a ‘tree prior’, typically a birth-death, Yule, or coalescent prior. Such nodes thus have the standard marginal prior and posterior distributions discussed above, and are conducive to the diptych metaphor. For those nodes that *are* directly calibrated using fossil information, the interpretation of the results of inference is more complicated. These nodes have a ‘user prior’, a distribution constructed in some way using information from the fossil record. However, these nodes are also involved in the tree prior. The resulting ‘marginal prior’ (or *effective* prior) is, in the BEAST implementation, a multiplicative combination of the user and tree priors. Furthermore, the marginal prior may also be influenced by interactions with adjacent user-calibrated nodes (e.g., ancestor and descendant nodes which have overlapping user priors), the complications of which are further exacerbated when topology is simultaneously inferred (see discussion in Rannala 2016). The resulting marginal prior thus does not necessarily reflect the original user prior. We note that this point has been raised previously in the methodological literature (Yang and Rannala 2006; Heled and Drummond 2012; Duchene *et al.* 2014; Heled and Drummond 2015; dos Reis 2016; Rannala 2016), and demonstrated empirically by Warnock *et al.* (2015), who report different behaviours in BEAST vs. MCMCtree (Yang 2007) implementations. Nevertheless, despite the fact that this is a recognized concern, the comparison of user and marginal priors is, in our judgment, *far too rarely assessed.* Finally, like uncalibrated nodes (and, indeed, all parameters in the model), fossil-calibrated nodes have marginal posterior distributions. However, unlike the uncalibrated nodes (which involve only two distributions, and thus a simple interpretation), calibrated nodes involve three distributions (two involving the prior, and one for the posterior), which complicates interpretation. [We prefer to continue with the diptych metaphor for calibrated nodes, rather than the obvious ‘triptych’, as the focus lies still on the change in belief on parameter values before (prior) and after (posterior) observing the data, even if the prior involves two components.]

For fossil-calibrated nodes, the difference between the marginal prior and marginal posterior, like the uncalibrated nodes, reflects information in the data (that is, the likelihood). However, the difference between the user and marginal priors, if present, may be better described as demonstrating the interaction of ‘pseudodata’ present in the various user priors. We justify this interpretation of pseudo-data in two ways. First, because divergence time estimation is only identifiable when incorporating temporal information, a prior on a node age is “more akin to a likelihood function than a prior” (Rannala 2016). Second, in the `BEAST` implementation of node-dating, each calibrated node has two priors (the user and tree priors); operationally, one of these serves the role of prior, while the other must serve as an elicited likelihood. Ideally, rather than being combined multiplicatively, the fossil calibration priors should be constructed so that they are conditional on the tree prior (Heled and Drummond 2012, 2015). An undetected interaction of priors, leading to a discrepancy between user and marginal priors, is especially concerning, as it would lead an investigator to believe the distinction between the user prior and marginal posterior is a result of information in the data (Rannala 2016). However, the interpretation of calibrations as pseudo-data holds regardless of whether there is an interaction that leads to a discrepancy from the original user priors.

We now turn our attention to the fossil-calibrated nodes in our empirical angiosperm example. Ideally we would find that the user and marginal priors are identical (that is, that the marginal priors reflect the intentions of the researcher), and the marginal priors and posteriors to differ (indicating information present in the data relevant to the parameter of interest). We plot in Fig. 3 the three sets of distributions for the 24 fossil-calibrated nodes for the Beaulieu *et al.* (2015) data set. In general, user and marginal priors match quite closely. However, there is a stark exception involving the *Tricolpites* constraint: while the user prior specifies that 95% of the prior mass should lie between 135.2-125 Ma, the marginal prior has a 95% highest posterior density (HPD) range of 226.1–128.7 Ma! The marginal posterior of this node age has a 95% HPD range of 170.7-146.6 Ma, which already surpasses the angiosperm paleontological estimate of ~140 Ma despite being well nested within the clade. We note that *Tricolpites* is the oldest constraint within angiosperms used by Beaulieu *et al.* (2015) (see Table 1 and Fig. 4). However, it is clear that it is not this particular calibration which is forcing angiosperms to be ‘too old’. Reanalysis without this specific constraint yielded even older posterior estimates for both this node (95% HPD: 181.2-154.3 Ma) and crown angiosperms (mean 241.0 Ma vs. 233.0 Ma with the constraint; data not shown). We cannot currently identify the cause of the disruption of the *Tricolpites* user prior.

**FIGURE 3.**
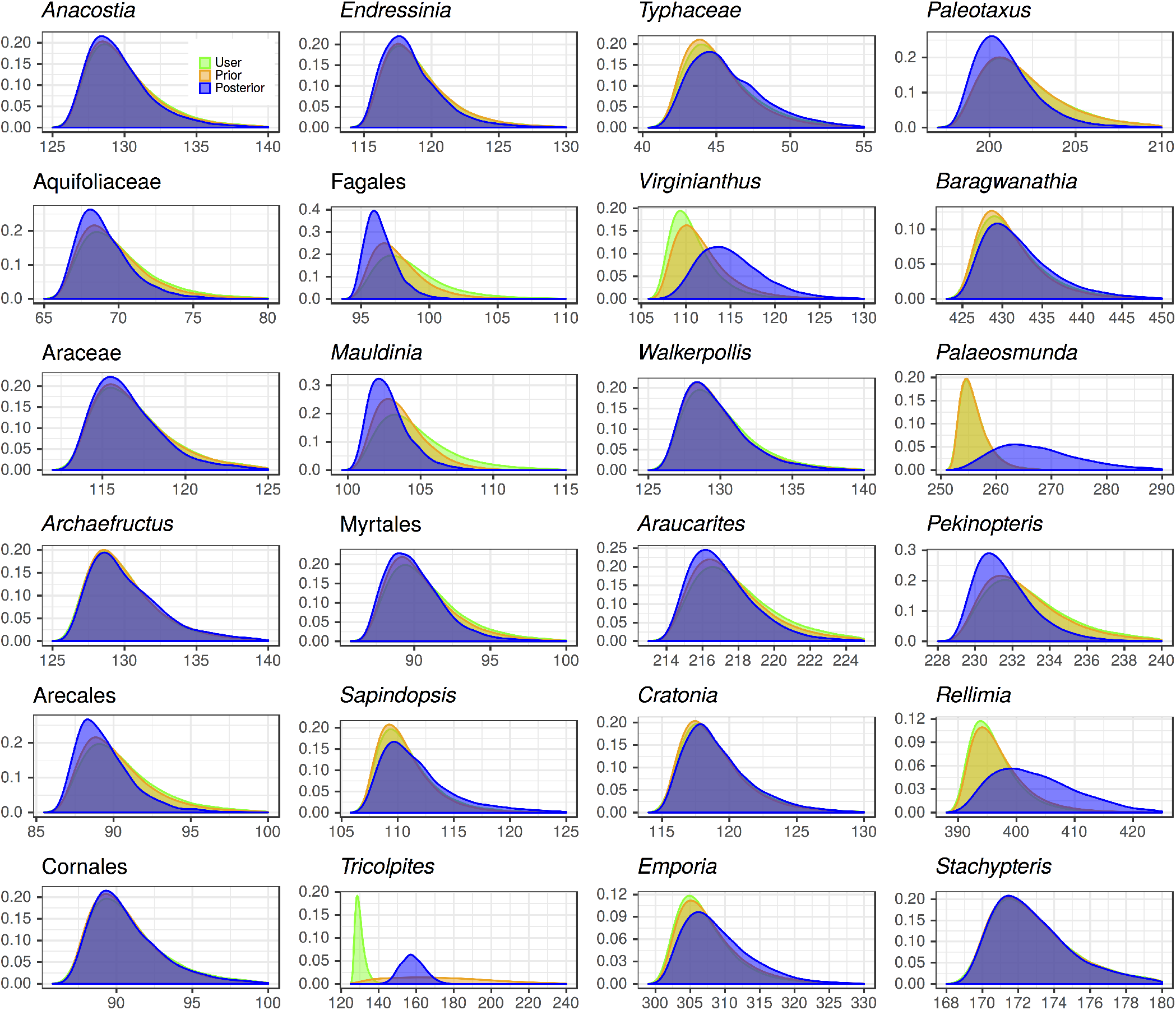
Densities for fossil-calibrated nodes from Beaulieu et al. (2015) (see Table 1). ‘User’ indicates the user-specified lognormal prior, ‘Prior’ indicates the marginal prior, and ‘Posterior’ indicates the marginal posterior. It is clear, for the majority of calibrated nodes, that (from the nearly complete overlap of prior and posterior distributions) the data contain insufficient information to overrule the priors for these parameters of interest.

Finally, we consider differences between the marginal prior and posterior distributions of age estimates for these same fossil-calibrated nodes. As discussed above, shifts in these paired distributions (i.e., following the diptych metaphor) would indicate the presence/degree of relevant phylogenetic ‘information’ (although theory has not yet been worked out on how to quantify this). However, from Fig. 3 we note that, for the most part, these distributions are nearly indistinguishable. This pattern is even more clear in Fig. 4 where, while uncalibrated nodes show significant shifts between prior and posterior analyses, calibrated nodes show essentially no movement. This lack of updating in belief from prior to posterior for the calibrated nodes suggests two non-mutually exclusive interpretations: (i) that the UCLN model, through allowing independent molecular substitution rates on all edges, may be overfitting the data, essentially allowing all calibrations to be mutually consistent (we do not expect, for example, that all of our calibration densities are equally ‘good’ – that is, that some constraints *should* conflict because of real vagaries of the fossil record); (ii) that the data set considered here lacks relevant phylogenetic ‘information’, or at least *insufficient* information to overrule the pseudo-data present in the fossil calibration densities.

**FIGURE 4.**
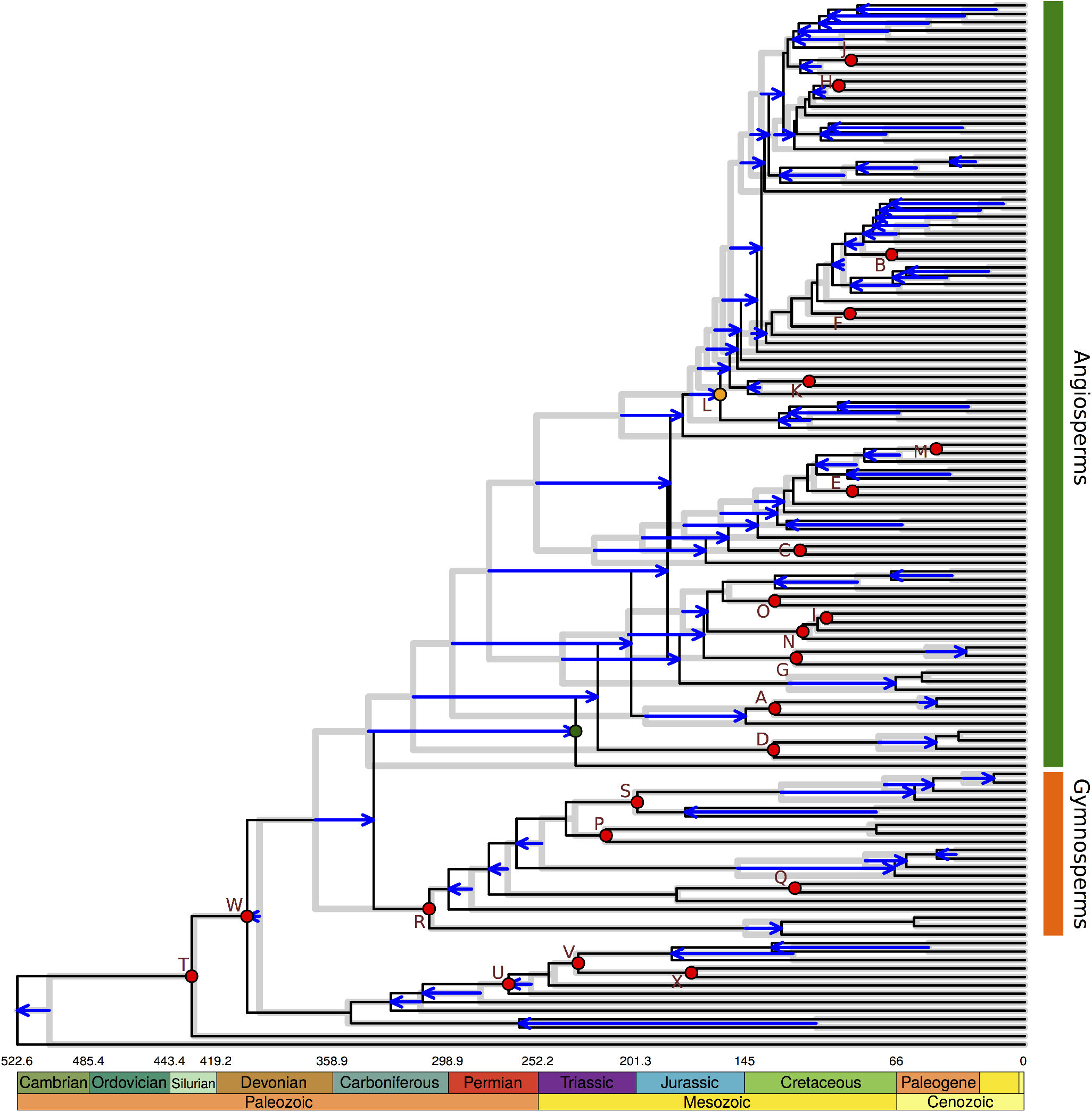
Mean prior (grey) and posterior (black) age estimates using the original lognormal calibration priors for the Beaulieu et al. (2015) data set. Arrows indicate shifts in age estimates from the prior to the posterior (nodes without arrows have shifted less than 5 Ma from prior to posterior). Nodes that are calibrated by fossil prior distributions are indicated with circles and letters (indicating fossil ids; see Table 1), except the *Tricolpites* node which is indicated with a yellow circle. Letters correspond to the fossil ids from Table 1. Finally, the (uncalibrated) crown angiosperm node is indicated by a green circle without a letter.

The finding of equivalent prior and posterior distributions may come as a surprise to some users, as unbounded (e.g., normal) or semi-bounded (e.g., lognormal) temporal priors are typically used to ‘let the data speak for themselves’. Certainly, divergence time inference is unusual in that it is only semi-identifiable (that is, only identifiable with both the sequence data *and* fossil calibrations; dos Reis and Yang 2013; Rannala 2016), so we do expect some level of association. It is, rather, the *degree* of association that is worrying. We are unaware of any other type of Bayesian analysis in evolutionary biology where identical prior and posterior distributions would not cause concern. The present results go a long way to explaining why divergence time estimation shows such a strong sensitivity to the fossil calibrations used (Inoue *et al.* 2010; Sauquet *et al.* 2012; Warnock *et al.* 2012; Warnock *et al.* 2015; Barba-Montoya *et al.* 2017; Warnock *et al.* 2017). The interpretation of fossil calibrations contributing pseudodata (rather than, say, fossils setting simple minimum age constraints as they have been traditionally) suggests that we might benefit from rethinking lessons that have been learned in the early days of phylogenetic divergence time estimation. We briefly consider one now.

## Are More Fossils Really Better In Node-Dating Analyses?

There is an adage in the divergence time estimation literature that as many fossils as possible should be used to calibrate nodes (Benton and Donoghue 2007). This makes sense, as of course we would like to include as much information as possible into a reconstruction. However, this advice largely came about when dating methods (e.g., r8s, Sanderson 2003; multidivtime, Thorne and Kishino 2002) employed constraints (e.g., Boolean minimum ages for the age of the fossil) rather than probabilistic distributions. As long as fossils were correctly placed within the phylogeny, the inclusion of more fossils should not produce misleading results. For instance, fossils that are ‘too young’ (that is, do not closely approximate in age the node they are calibrating) are either simply uninformative, or appropriately represent limitations of the fossil record. As an extreme example, a chicken bone found in a back alley gutter is a valid (if imprecise) minimum age constraint for the origin of *Gallus gallus* (recently estimated at 2.9 Ma; Stein *et al.* 2015).

However, from the results reported above (Figs. 3,4) we find several concerns with including as many fossils as possible in node-dating analyses that employ routine parametric (e.g., lognormal) fossil calibration distribution priors. [We note that these concerns do not apply to the fossilized birth-death model (Heath *et al.* 2014) or tip-dating (Ronquist *et al.* 2012) approaches to divergence time estimation, which do not involve such calibrations.] First, as with the *Tricolpites* example above, calibrations can interact with each other and the tree prior in unpredictable ways to produce marginal priors that do not represent the originally intended user priors. While this is a recognized (though under-appreciated) issue, the available solutions in the `BEAST` implementation work only for a small number of calibrations (Heled and Drummond 2012; Heled and Drummond 2015), and Rannala (2016) demonstrates that a general solution to preserving user calibration priors when jointly inferring topology is not possible. Second, given that the marginal prior and posterior calibrated node ages are often indistinguishable (suggesting little relevant phylogenetic information content), it is worrisome that the act of employing such temporal calibration priors can directly determine the resulting posterior patterns of rate heterogeneity across a tree from signal in the prior (rather than from signal in the data). Any rate estimates that arise from such an analysis should be regarded with skepticism. It is not inconceivable, for example, that the parametric use of the best available fossils from an incomplete fossil record could turn a clock-like data set unclocklike, needlessly increasing the model complexity (and therefore uncertainty) involved.

Our final concern with unrestrained parametric calibration use is the form of the calibrations themselves. A flexible assortment of distribution families are available (Ho and Phillips 2009; see also discussion in Brown and van Tuinen 2011), allowing essentially any prior belief to be employed. In addition, researchers can make use of the fossil calibration database (Ksepka *et al.* 2015), and prescribed ‘best practices’ (Parham *et al.* 2012) can help avoid naïve errors when dealing with the fossil record. Nevertheless, the vast majority of user calibration priors employed in the literature are wholly idiosyncratic and arbitrary (we include ourselves here). This is not necessarily a result of molecular phylogeneticists lacking the appropriate paleontological expertise (and isn’t that what collaboration is for?), but rather a property of data involved.

While methods exist to generate a distribution from a set of fossils (Marshall 2008; Nowak *et al.* 2013; Claramunt and Cracraft 2015), these require well-sampled data Scant data is an entirely different problem. How does one fit a distribution to a single (exceptionally old, and therefore exceptionally informative) fossil? Minimum bounds are simple (the age of the oldest fossil), but as Parham *et al.* (2012) note, there exists no standard protocol for formulating maximum ages (let alone the shape of the distribution spanning the upper and lower bounds). [A newly developed method by Matschiner *et al.* (2017) is intuitively and empirically promising, but requires a researcher to supply rates of speciation, extinction, and fossilization, reliable estimates of which might not be available for a focal lineage.] Indeed, the process of constructing temporal priors is so nebulous that Lee and Skinner (2011) likened it to “educated guesswork”. However, it is not the arbitrariness of the calibrations *per se* that is of concern, but rather that they act as a strong source of pseudo-data Taken to a hyperbolic extreme, if calibration priors were applied to every node in a tree, then the results above would suggest that there would be no use in running the analysis at all. More worrisome is that such a chronogram may largely determine the results of downstream comparative analyses involving either quantifying lineage diversification rate heterogeneity (Rabosky, 2014) or inferring ancestral character states (Sauquet *et al.*, 2017). In other types of Bayesian analysis we expect that an increase in data can overrule poorly constructed priors and converge on an answer. It is presently unclear whether this is true when performing node-dating using the UCLN model and routinely constructed lognormal fossil calibration priors, and if so, how much data (or ‘information’, whatever that turns out to be) would be required. It is well known that the uncertainty in divergence time estimates cannot be reduced arbitrarily, even with infinite amounts of data (Yang and Rannala 2006; Rannala and Yang 2007; dos Reis and Yang 2013), but it is unclear how data can override the pseudo-data present in the node calibration priors.

## Can Uniform Calibration Priors Solve Issues In Node-Dating?

A concern expressed above is that the fossil prior calibration distributions typically employed in node-dating analyses, through their role as pseudo-data, may unduly (and inadvertently) restrict the breadth of posterior parameter values. A possible remedy to this problem may therefore be to relax the excessive influence of fossil calibration distributions through employing them instead as extremely broad uniform priors. This approach has been applied previously (e.g. Tong *et al.* 2015; Foster *et al.* 2017). We therefore explored the influence of this type of approach on the Beaulieu *et al.* (2015) data set by specifying broad uniform calibration priors to all Spermatophyta (angiosperm and gymnosperm) fossil-calibrated nodes. For these nodes, the minimum of the uniform distribution was set to the age of the fossil (Table 1), while the maximum was set to 350 Ma as recommended by Magallón and Castillo (2009) (and implemented for some nodes in Foster *et al.* 2017). It is not possible to objectively identify optimal maximum values for these distributions, so the value of 350 Ma should be regarded as useful for illustrative purposes. Non-spermatophyte fossils were calibrated with the original lognormal distributions (Table 1), and BEAST MCMC settings are the same as above.

The goal of employing broad uniform fossil calibration priors is to diffuse the prior mass sufficiently such that the tree (e.g., birth-death) prior dominates in the prior calculations. Collectively, the minimum node age bounds (i.e., the fossil ages) combine with the tree prior to form a joint prior which is effectively a truncated birth-death prior. Results from these analyses show this to be the case (Fig. 5), as the strict recapitulation of the prior by the posterior is largely eliminated. The marginal node priors produced therefore reflect the joint truncated birth-death prior.

**FIGURE 5.**
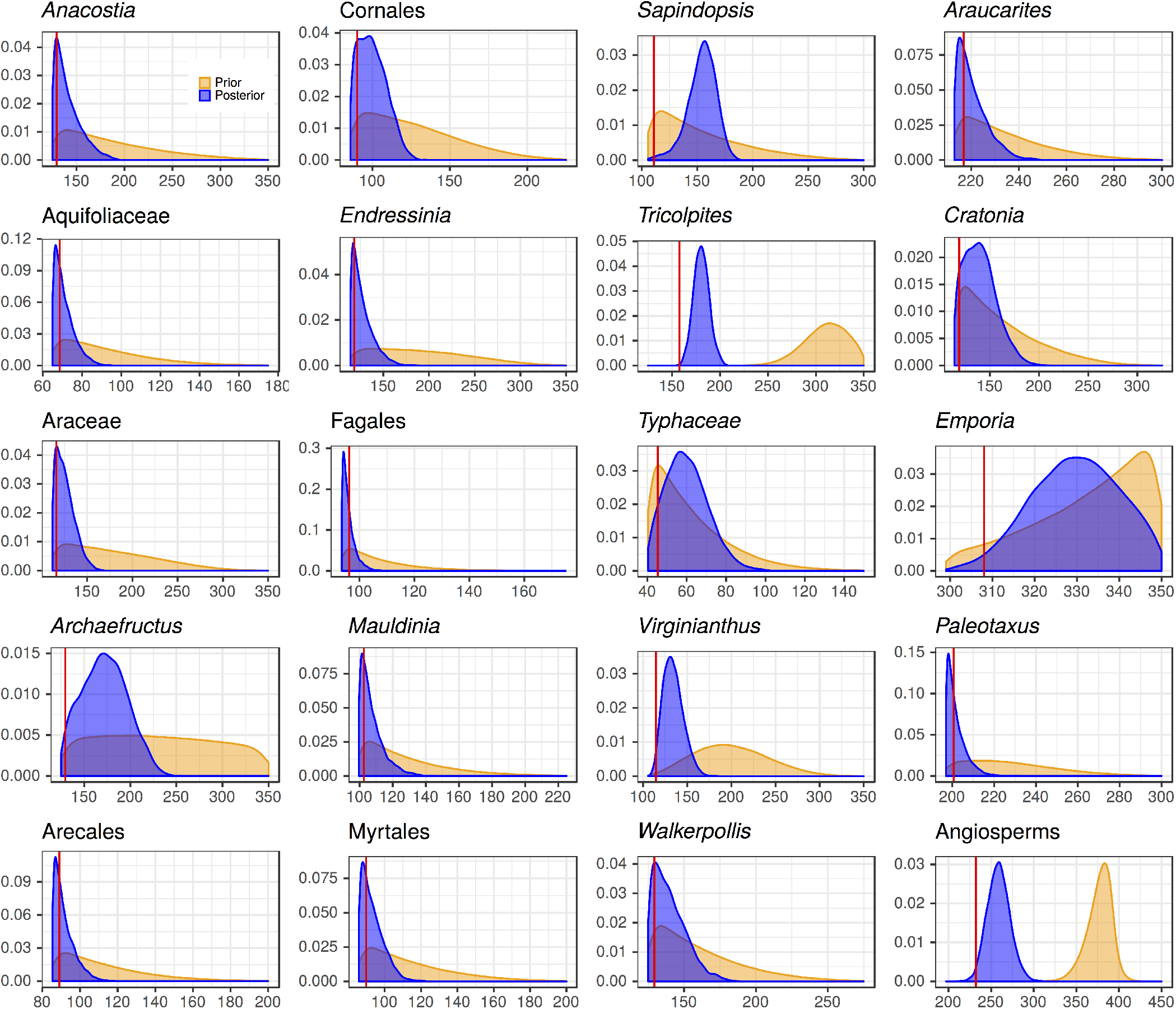
Densities for fossil-calibrated spermatophyte nodes from Beaulieu et al. (2015) using broad uniform fossil calibration priors with maximum bounds set to 350 Ma (see text for details). Note that calibrated non-spermatophyte nodes are not shown as they employ lognormal calibration priors and thus do not differ from the previous results. As above, ‘Prior’ indicates the marginal prior, and ‘Posterior’ indicates the marginal posterior. Red vertical lines indicate the median posterior estimate using lognormal calibration priors (see Fig. 3). As in the analyses above, the crown angiosperms node is not calibrated directly by a fossil.

So, are diffuse uniform fossil calibration priors the solution to concerns in node-dating divergence time estimation? There are some clear positive aspects: (i) they can remove the strict correspondence between prior and posterior parameter distributions seen with lognormal calibration priors (Fig. 3), and (ii) they attempt to use the fossil record at face value (i.e., through employing hard minimums set to fossil ages). However, we also note some additional concerns. First, the use of broad uniform priors appears somewhat disingenuous. For example, the mean uniform fossil prior range for Foster *et al.* (2017) was 116.9 Ma (median 82.4 Ma); researchers are unlikely to *genuinely* have a prior belief that a divergence time is equally probable over a 100 Ma period. Rather, such priors are employed in an ad hoc manner simply to enable the analysis. Although the lognormal priors discussed above are quite arbitrary, they are at least genuine in their attempt to approximate actual prior belief in the correspondence fossil ages and actual divergence times. Second, on a related issue, the choice of an explicit (hard) upper bound (as discussed above) is predominantly subjective. Although outside of the scope of the present study, an upper bound on the age of a node clearly restricts prior (and hence, posterior) parameter estimates, potentially with misleadingly high precision (Yang and Rannala, 2006). Finally, the results from Fig. 5 show the influence of the joint truncated birth-death prior. It is clear that, within the confines of the joint prior, several age constraints conflict strongly with one another. For example, the *Emporia* and *Tricolpites* constraints collide with the upper bound, suggesting they would accommodate older ages if permitted. At the other extreme, several priors appear to pile up against the minimum boundary. Taken together, these results suggest that the tree prior (here, a birth-death prior) is not consistent with the entire set of fossil ages. This suggests that a homogeneous birth-death prior may be ill-suited to these data. This result is reminiscent of the influence of the extent of taxonomic sampling on inferred divergence time estimates from Beaulieu *et al.* (2015). Thus, if it is understood from other means that a homogeneous birth-death process is unlikely to adequately describe an entire tree, it may be preferable to explicitly anchor nodes according to some justifiable distribution (e.g., Claramunt and Cracraft 2015).

## Where To Go From Here?

The results presented here highlight several issues that should be considered as the field moves forward. In regard to angiosperms, is the amount of temporal ‘information’ present in the molecular data (or, on the other hand, the adequacy of current relaxed clock models) insufficient to reconstruct such recalcitrant nodes as the age of crown angiosperms? If this is the case, the only way forward, given the methods of inference, may be to apply user priors that are *intended* to constitute pseudo-data. However, if prudent, this fact needs to be more widely recognized and acknowledged. In some respects this is a defensible position as, if fossil calibrations are constructed with significant information about the fossil record, estimations will be constrained *to* existing fossil information. In this vein, the results of Magallón *et al.* (2015), which estimate nested angiosperm divergence times within a strict paleontologically-imposed age of the ancestral crown node, are reasonable in the context of the data available (Sanderson 2015). Nevertheless, it should be made clear when the molecular data, in this context, do not significantly alter the posterior distribution. If this is indeed the way forward, then care should be taken to assess both the validity of the fossils being used (Sanders and Lee 2007; Brown and Sorhannus 2010) and the form of the calibration priors (Inoue *et al.* 2010; Brown and Sorhannus 2010; Sauquet *et al.* 2012; Warnock *et al.* 2012; Duchene *et al.* 2014; Barba-Montoya *et al.* 2017; Foster *et al.* 2017) Regarding the latter, we note that constructing calibration distributions using hyperpriors (Heath, 2012) or an explicit modelling of fossil sampling probabilities (Silvestro *et al.* 2015; Matschiner *et al.* 2017; Warnock *et al.* 2017) are attractive possibilities. Nevertheless, we strongly advocate the regular use of the diptych approach to data analysis by habitually comparing prior and posterior distributions: it is imperative to understand which parameters in our models are informed by the data present, and which simply recapitulate the prior. When hypothesis testing it is even more critically important to determine whether a hypothesis is rejected by the data or, as with the crown angiosperm age results above, are effectively precluded by the joint prior.

However, new methods of divergence time inference are emerging that largely bypass the concerns associated with node-dating (see reviews in Heath and Moore 2014; Donoghue and Yang 2016). The fossilized birth-death model of Heath *et al.* (2014) incorporates extant and extinct (i.e., sampled fossils) lineages as evolving according to the same underlying diversification model. Alternatively, when morphological data are available for both extinct and extant taxa, divergence times can be estimated using the tip-dating approach of Ronquist *et al.* (2012). Both of these methods, and combinations thereof (Zhang *et al.* 2015; Gavryushkina *et al.* 2017) can take advantage of an arbitrary number of fossils within a lineage (rather than being reduced to a single distribution as in node-dating) and incorporate fossil temporal information directly without extrapolation. The excitement surrounding these methods might lead us to think it not unreasonable to suppose that in the near future node-dating will be regarded as a useful tool that was ultimately replaced by methods that more directly make use of the available data. However, both of these methods are relatively new, and it is unclear whether they will overthrow node-dating results for the most recalcitrant nodes (i.e., placental mammals, crown birds, crown angiosperms, etc.). These methods also raise new questions in regard to model adequacy, implied and explicit assumptions regarding both diversification and morphology models, and data availability and quality for extinct and extant lineages. Furthermore, the resulting divergence time estimates from these new methods may not differ as much as expected. For example, Eguchi and Tamura (2016) employed the fossilized birth-death model and found monocots arose 174.26-134.14 Ma, which does not conflict strongly with previous node-dating results.

## Final Thoughts On The Age Of Angiosperms

Finally, we note that the results presented above do not refute the concerns identified and demonstrated through simulation by Beaulieu *et al.* (2015) regarding violations of biological modelling. While among-lineage molecular substitution rate heterogeneity is regarded as ubiquitous, clade- and trait-specific correlations of rate variation explored by Beaulieu *et al.* (2015) are becoming increasingly recognized as important biological patterns of molecular evolution (Smith *et al.* 2010; Dornburg *et al.* 2012; Lartillot and Delsuc 2012; Worobey *et al.* 2014). Such processes must be correctly modelled if our divergence time estimates are to be accurate. In this vein, we note that the fit (Lepage *et al.* 2007; Ho *et al.* 2015) and adequacy (Duchene *et al.*, 2015) of alternative clock models are far too rarely assessed. In addition, the artefacts of lineage sampling identified by Beaulieu *et al.* (2015) casts doubt on the suitability of a homogeneous birth-death model as a prior on node ages. This doubt is especially manifested with respect to dating the evolution of angiosperms, where it is known *a priori* that lineages exhibit an incredible breadth of diversification rates (Tank *et al.*, 2015), not to mention that the extant angiosperm diversity dwarfs other embryophyte clades. We thus regard our results above as complementary to those of Beaulieu *et al.* (2015), and only with both in mind can we confidently move forward. Although provocative, we hope that our discussion above will help cultivate a deeper conversation on how best to proceed.

## Supplementary Material

Data available from the Dryad Data Repository: http://dx.doi.org/10.5061/dryad.f20j0.

## Acknowledgements

We thank Jeremy Beaulieu and Brian O’Meara for sharing their thoughts on these issues, and Susana Magallóon for graciously sharing her BEAST xml input file. We would also like to thank the following for productive discussions (although none necessarily hold the views expressed herein): Ben Redelings, Joseph Walker, Simon Uribe-Convers, Caroline Parins-Fukuchi, Ning Wang, Greg Stull, Aaron King, Michael Landis, Michael Matschiner, Alex Taylor, Oscar Vargas, Paul Lewis, and members of the Smith laboratory. We would like to explicitly and emphatically thank Hervóe Sauquet for providing a thoughtful review of the bioRxiv preprint version of this manuscript. Associate Editor Simon Ho provided invaluable feedback, both within and outside of the recognized review process, which helped to improve the manuscript. J.W.B. thanks Don Van Vliet for help with the title, Emo Philips for help with broader dissemination, Mark Kozelek for encouragement, and Linda Slote for tracking down an obscure Testudines reference (that was unfortunately dropped from an earlier version of this paper).

## REFERENCES

Ayres, D. L., Darling, A., Zwickl, D. J., Beerli, P., Holder, M. T., Lewis, P. O., Huelsenbeck, J. P., Ronquist, F., Swofford, D. L., Cummings, M. P., Rambaut, A., and Suchard, M. A. 2012. BEAGLE: an application programming interface and high-performance computing library for statistical phylogenetics. Systematic Biology, 61(1): 170–173.

Barba-Montoya, J., dos Reis, M., and Yang, Z. 2017. Comparison of different strategies for using fossil calibrations to generate the time prior in bayesian molecular clock dating. Molecular Phylogenetics and Evolution.

Beaulieu, J. M., O’Meara, B. C., Crane, P., and Donoghue, M. J. 2015. Heterogeneous rates of molecular evolution and diversification could explain the Triassic age estimate for angiosperms. Systematic Biology, 64(5): 869–878.

Bell, C. D., Soltis, D. E., and Soltis, P. S. 2010. The age and diversification of the angiosperms re-revisited. American Journal of Botany, 97(8): 1296–1303.

Benton, M. J. 1999. Early origins of modern birds and mammals: molecules vs. morphology. BioEssays, 21(12): 1043–1051.

Benton, M. J. and Donoghue, P. C. J. 2007. Paleontological evidence to date the tree of life. Molecular Biology and Evolution, 24(1): 26–53.

Brenner, G. J. 1996. Evidence for the earliest stage of angiosperm pollen evolution: A paleoequatorial section from israel. In D. W. Taylor and L. J. Hickey, editors, Flowering Plant Origin, Evolution & Phylogeny, pages 91–115. Springer US, Boston, MA.

Brown, J. W. and Sorhannus, U. 2010. A molecular genetic timescale for the diversification of autotrophic stramenopiles (Ochrophyta): Substantive underestimation of putative fossil ages. PLoS ONE, 5(9): 1–11.

Brown, J. W. and van Tuinen, M. 2011. Evolving perceptions on the antiquity of the modern avian tree. In G. Dyke and G. Kaiser, editors, Living Dinosaurs: The Evolutionary History of Modern Birds, chapter 12, pages 306–324. John Wiley & Sons Ltd, Oxford.

Brown, J. W., Payne, R. B., and Mindell, D. P. 2007. Nuclear DNA does not reconcile ‘rocks’ and ‘clocks’ in Neoaves: a comment on Ericson et al. Biology Letters, 3(3): 257–260.

Brown, J. W., Walker, J. F., and Smith, S. A. 2017. Phyx: phylogenetic tools for unix. Bioinformatics, 33(12): 1886–1888.

Claramunt, S. and Cracraft, J. 2015. A new time tree reveals Earth history’s imprint on the evolution of modern birds. Science Advances, 1(11): e1501005.

Donoghue, P. C. J. and Benton, M. J. 2007. Rocks and clocks: calibrating the tree of life using fossils and molecules. Trends in Ecology & Evolution, 22(8): 424–431.

Donoghue, P. C. J. and Yang, Z. 2016. The evolution of methods for establishing evolutionary timescales. Philosophical Transactions of the Royal Society of London B: Biological Sciences, 371(1699): 20160020.

Dornburg, A., Brandley, M. C., McGowen, M. R., and Near, T. J. 2012. Relaxed clocks and inferences of heterogeneous patterns of nucleotide substitution and divergence time estimates across whales and dolphins (Mammalia: Cetacea). Molecular Biology and Evolution, 29(2): 721–736.

dos Reis, M. 2016. Notes on the birth-death prior with fossil calibrations for Bayesian estimation of species divergence times. Philosophical Transactions of the Royal Society of London B: Biological Sciences, 371(1699): 20150128.

dos Reis, M. and Yang, Z. 2013. The unbearable uncertainty of Bayesian divergence time estimation. Journal of Systematics and Evolution, 51(1): 30–43.

Drummond, A. J. and Rambaut, A. 2007. BEAST: Bayesian evolutionary analysis by sampling trees. BMC Evolutionary Biology, 7(1): 214.

Drummond, A. J., Ho, S. Y. W., Phillips, M. J., and Rambaut, A. 2006. Relaxed phylogenetics and dating with confidence. PLoS Biology, 4(5).

Duchene, D. A., Duchene, S., Holmes, E. C., and Ho, S. Y. W. 2015. Evaluating the adequacy of molecular clock models using posterior predictive simulations. Molecular Biology and Evolution, 32(11): 2986–2995.

Duchene, S. and Ho, S. Y. W. 2014. Using multiple relaxed-clock models to estimate evolutionary timescales from DNA sequence data. Molecular Phylogenetics and Evolution, 77: 65–70.

Duchene, S., Lanfear, R., and Ho, S. Y. W. 2014. The impact of calibration and clock-model choice on molecular estimates of divergence times. Molecular Phylogenetics and Evolution, 78: 277–289.

Eguchi, S. and Tamura, M. N. 2016. Evolutionary timescale of monocots determined by the fossilized birth-death model using a large number of fossil records. Evolution, 70(5): 1136–1144.

Ericson, P. G., Anderson, C. L., Britton, T., Elzanowski, A., Johansson, U. S., Källersjö, M., Ohlson, J. I., Parsons, T. J., Zuccon, D., and Mayr, G. 2006. Diversification of Neoaves: integration of molecular sequence data and fossils. Biology Letters, 2(4): 543–547.

Foster, C. S. P., Sauquet, H., van der Merwe, M., McPherson, H., Rossetto, M., and Ho, S. Y. W. 2017. Evaluating the impact of genomic data and priors on Bayesian estimates of the angiosperm evolutionary timescale. Systematic Biology, 66(3): 338–351.

Gavryushkina, A., Heath, T. A., Ksepka, D. T., Stadler, T., Welch, D., and Drummond, A. J. 2017. Bayesian total-evidence dating reveals the recent crown radiation of penguins. Systematic biology, 66(1): 57–73.

Heath, T. A. 2012. A hierarchical Bayesian model for calibrating estimates of species divergence times. Systematic Biology, 61(5): 793–809.

Heath, T. A. and Moore, B. R. 2014. Bayesian inference of species divergence times. In M.-H. Chen, L. Kuo, and P. O. Lewis, editors, Bayesian Phylogenetics: Methods Algorithms, and Applications, chapter 13, pages 277–318. CRC Press, Boca Raton, Florida.

Heath, T. A., Huelsenbeck, J. P., and Stadler, T. 2014. The fossilized birthdeath process for coherent calibration of divergence-time estimates. Proceedings of the National Academy of Sciences, 111(29): E2957–E2966.

Heibl, C. 2008. Phyloch: R language tree plotting tools and interfaces to diverse phylogenetic software packages.

Heled, J. and Drummond, A. J. 2012. Calibrated tree priors for relaxed phylogenetics and divergence time estimation. Systematic Biology, 61(1): 138–149.

Heled, J. and Drummond, A. J. 2015. Calibrated birth-death phylogenetic time-tree priors for Bayesian inference. Systematic Biology, 64(3): 369–383.

Herendeen, P. S., Friis, E. M., Pedersen, K. R., and Crane, P. R. 2017. Palaeobotanical redux: revisiting the age of the angiosperms. Nature Plants, 3: 17015.

Ho, S. Y. W. and Phillips, M. J. 2009. Accounting for calibration uncertainty in phylogenetic estimation of evolutionary divergence times. Systematic Biology, 58(3): 367–380.

Ho, S. Y. W., Duchene, S., and Duchene, D. 2015. Simulating and detecting autocorrelation of molecular evolutionary rates among lineages. Molecular Ecology Resources, 15(4): 688–696.

Holland, S. M. 2016. The non-uniformity of fossil preservation. Philosophical Transactions of the Royal Society of London B: Biological Sciences, 371(1699): 20150130.

Inoue, J., Donoghue, P. C. J., and Yang, Z. 2010. The impact of the representation of fossil calibrations on Bayesian estimation of species divergence times. Systematic Biology, 59(1): 74–89.

Jarvis, E. D., Mirarab, S., Aberer, A. J., Li, B., Houde, P., Li, C., Ho, S. Y. W., Faircloth, B. C., Nabholz, B., Howard, J. T., Suh, A., Weber, C. C., da Fonseca, R. R., Li, J., Zhang, F., Li, H., Zhou, L., Narula, N., Liu, L., Ganapathy, G., Boussau, B., Bayzid, M. S., Zavidovych, V., Subramanian, S., Gabaldón, T., Capella-Gutierrez, S., Huerta-Cepas, J., Rekepalli, B., Munch, K., Schierup, M., Lindow, B., Warren, W. C., Ray, D., Green, R. E., Bruford, M. W., Zhan, X., Dixon, A., Li, S., Li, N., Huang, Y., Derryberry, E. P., Bertelsen, M. F., Sheldon, F. H., Brumfield, R. T., Mello, C. V., Lovell, P. V., Wirthlin, M., Schneider, M. P. C., Prosdocimi, F., Samaniego, J. A., Velazquez, A. M. V., Alfaro-Nóñez, A., Campos, P. F., Petersen, B., Sicheritz-Ponten, T., Pas, A., Bailey, T., Scofield, P., Bunce, M., Lambert, D. M., Zhou, Q., Perelman, P., Driskell, A. C., Shapiro, B., Xiong, Z., Zeng, Y., Liu, S., Li, Z., Liu, B., Wu, K., Xiao, J., Yinqi, X., Zheng, Q., Zhang, Y., Yang, H., Wang, J., Smeds, L., Rheindt, F. E., Braun, M., Fjeldsa, J., Orlando, L., Barker, F. K., Jønsson, K. A., Johnson, W., Koepfli, K.-P., O’Brien, S., Haussler, D., Ryder, O. A., Rahbek, C., Willerslev, E., Graves, G. R., Glenn, T. C., McCormack, J., Burt, D., Ellegren, H., Alstrom, P., Edwards, S. V., Stamatakis, A., Mindell, D. P., Cracraft, J., Braun, E. L., Warnow, T., Jun, W., Gilbert, M. T. P., and Zhang, G. 2014. Whole-genome analyses resolve early branches in the tree of life of modern birds. Science, 346(6215): 1320–1331.

Ksepka, D. T., Ware, J. L., and Lamm, K. S. 2014. Flying rocks and flying clocks: disparity in fossil and molecular dates for birds. Proceedings of the Royal Society of London B: Biological Sciences, 281(1788): 20140677.

Ksepka, D. T., Parham, J. F., Allman, J. F., Benton, M. J., Carrano, M. T., Cranston, K. A., Donoghue, P. C. J., Head, J. J., Hermsen, E. J., Irmis, R. B., Joyce, W. G., Kohli, M., Lamm, K. D., Leehr, D., Patan, J. L., Polly, P. D., Phillips, M. J., Smith, N. A., Smith, N. D., Van Tuinen, M., Ware, J. L., and Warnock, R. C. M. 2015. The fossil calibration database-a new resource for divergence dating. Systematic Biology, 64(5): 853–859.

Lartillot, N. and Delsuc, F. 2012. Joint reconstruction of divergence times and lifehistory evolution in placental mammals using a phylogenetic covariance modell. Evolution, 66(6): 1773–1787.

Lee, M. S. Y. and Skinner, A. 2011. Testing fossil calibrations for vertebrate molecular trees. Zoologica Scripta, 40(5): 538–543.

Lepage, T., Bryant, D., Philippe, H., and Lartillot, N. 2007. A general comparison of relaxed molecular clock models. Molecular Biology and Evolution, 24(12): 2669–2680.

Lewis, P. O., Chen, M.-H., Kuo, L., Lewis, L. A., Fučíkovó, K., Neupane, S., Wang, Y.-B., and Shi, D. 2016. Estimating Bayesian hylogenetic information content. Systematic Biology, 65(6): 1009–1023.

Magallón, S. and Castillo, A. 2009. Angiosperm diversification through time. American Journal of Botany, 96(1): 349–365.

Magallón, S., Hilu, K. W., and Quandt, D. 2013. Land plant evolutionary timeline: Gene effects are secondary to fossil constraints in relaxed clock estimation of age and substitution rates. American Journal of Botany, 100(3): 556–573.

Magallón, S., Goómez-Acevedo, S., Saónchez-Reyes, L. L., and Hernaóndez-Hernóandez, T. 2015. A metacalibrated time-tree documents the early rise of flowering plant phylogenetic diversity. New Phytologist, 207(2): 437–453.

Marshall, C. R. 2008. A simple method for bracketing absolute divergence times on molecular phylogenies using multiple fossil calibration points. The American Naturalist, 171(6): 726–742.

Matschiner, M., Musilova, Z., Barth, J. M. I., Starostovó, Z., Salzburger, W., Steel, M., and Bouckaert, R. 2017. Bayesian phylogenetic estimation of clade ages supports trans-atlantic dispersal of cichlid fishes. Systematic Biology, 66(1): 3–22.

Meredith, R. W., Janecka, J. E., Gatesy, J., Ryder, O. A., Fisher, C. A., Teeling, E. C., Goodbla, A., Eizirik, E., Simñao, T. L. L., Stadler, T., Rabosky, D. L., Honeycutt, R. L., Flynn, J. J., Ingram, C. M., Steiner, C., Williams, T. L., Robinson, T. J., Burk-Herrick, A., Westerman, M., Ayoub, N. A., Springer, M. S., and Murphy, W. J. 2011. Impacts of the Cretaceous terrestrial revolution and KPg extinction on mammal diversification. Science, 334(6055): 521–524.

Mitchell, K. J., Cooper, A., and Phillips, M. J. 2015. Comment on “Whole-genome analyses resolve early branches in the tree of life of modern birds”. Science, 349(6255): 1460–1460.

Nowak, M. D., Smith, A. B., Simpson, C., and Zwickl, D. J. 2013. A simple method for estimating informative node age priors for the fossil calibration of molecular divergence time analyses. PLoS ONE, 8(6): 1–13.

O’Leary, M. A., Bloch, J. I., Flynn, J. J., Gaudin, T. J., Giallombardo, A., Giannini, N. P., Goldberg, S. L., Kraatz, B. P., Luo, Z.-X., Meng, J., Ni, X., Novacek, M. J., Perini, F. A., Randall, Z. S., Rougier, G. W., Sargis, E. J., Silcox, M. T., Simmons, N. B., Spaulding, M., Velazco, P. M., Weksler, M., Wible, J. R., and Cirranello, A. L. 2013. The placental mammal ancestor and the post-K-Pg radiation of placentals. Science, 339(6120): 662–667.

Parham, J. F., Donoghue, P. C. J., Bell, C. J., Calway, T. D., Head, J. J., Holroyd, P. A., Inoue, J. G., Irmis, R. B., Joyce, W. G., Ksepka, D. T., Patane, J. S. L., Smith, N. D., Tarver, J. E., van Tuinen, M., Yang, Z., Angielczyk, K. D., Greenwood, J. M., Hipsley, C. A., Jacobs, L., Makovicky, P. J., Mller, J., Smith, K. T., Theodor, J. M., Warnock, R. C. M., and Benton, M. J. 2012. Best practices for justifying fossil calibrations. Systematic Biology, 61(2): 346–359.

Phillips, M. J. 2009. Branch-length estimation bias misleads molecular dating for a vertebrate mitochondrial phylogeny. Gene, 441(12): 132–140.

Prum, R. O., Berv, J. S., Dornburg, A., Field, D. J., Townsend, J. P., Lemmon, E. M., and Lemmon, A. R. 2015. A comprehensive phylogeny of birds (Aves) using targeted next-generation DnA sequencing. Nature, 526(7574): 569–573.

R Core Team 2016. R: A Language and Environment for Statistical Computing. R Foundation for Statistical Computing, Vienna, Austria.

Rabosky, D. L. 2014. Automatic detection of key innovations, rate shifts, and diversity-dependence on phylogenetic trees. PloS one, 9(2): e89543.

Rannala, B. 2016. Conceptual issues in Bayesian divergence time estimation. Philosophical Transactions of the Royal Society of London B: Biological Sciences, 371(1699): 20150134.

Rannala, B. and Yang, Z. 2007. Inferring speciation times under an episodic molecular clock. Systematic Biology, 56(3): 453–466.

Revell, L. J., Harmon, L. J., Glor, R. E., and Linder, P. 2005. Under-parameterized model of sequence evolution leads to bias in the estimation of diversification rates from molecular phylogenies. Systematic Biology, 54(6): 973–983.

Ronquist, F., Klopfstein, S., Vilhelmsen, L., Schulmeister, S., Murray, D. L., and Rasnitsyn, A. P. 2012. A total-evidence approach to dating with fossils, applied to the early radiation of the Hymenoptera. Systematic Biology, 61(6): 973–999.

Sanders, K. L. and Lee, M. S. 2007. Evaluating molecular clock calibrations using Bayesian analyses with soft and hard bounds. Biology Letters, 3(3): 275–279.

Sanderson, M. J. 2003. r8s: inferring absolute rates of molecular evolution and divergence times in the absence of a molecular clock. Bioinformatics, 19(2): 301–302.

Sanderson, M. J. 2015. Back to the past: a new take on the timing of flowering plant diversification. New Phytologist, 207(2): 257259.

Sauquet, H., Ho, S. Y. W., Gandolfo, M. A., Jordan, G. J., Wilf, P., Cantrill, D. J., Bayly, M. J., Bromham, L., Brown, G. K., Carpenter, R. J., Lee, D. M., Murphy, D. J., Sniderman, J. M. K., and Udovicic, F. 2012. Testing the impact of calibration on molecular divergence times using a fossil-rich group: The case of Nothofagus (Fagales). Systematic Biology, 61(2): 289–313.

Sauquet, H., von Balthazar, M., Magallón, S., Doyle, J. A., Endress, P. K., Bailes, E. J., Morais, E. B. d., Bull-Hereñu, K., Carrive, L., Chartier, M., Chomicki, G., Coiro, M., Cornette, R., El Ottra, J. H. L., Epicoco, C., Foster, C. S. P., Jabbour, F., Haevermans, A., Haevermans, T., Hernóndez, R., Little, S. A., Löfstrand, S., Luna, J. A., Massoni, J., Nadot, S., Pamperl, S., Prieu, C., Reyes, E., Santos, P. d., Schoonderwoerd, K. M., Sontag, S., Soulebeau, A., Staedler, Y., Tschan, G. F., Leung, A. W.-S., and Schönenberger, J. 2017. The ancestral flower of angiosperms and its early diversification. Nature Communications, 8: ncomms16047.

Schenk, J. J. and Hufford, L. 2010. Effects of substitution models on divergence time estimates: Simulations and an empirical study of model uncertainty using cornales. Systematic Botany, 35(3): 578–592.

Shannon, C. E. 1948. Modeling compositional heterogeneity. Bell System Technical Journal, 27: 379–423, 623-656.

Silvestro, D., Cascales-Minñana, B., Bacon, C. D., and Antonelli, A. 2015. Revisiting the origin and diversification of vascular plants through a comprehensive Bayesian analysis of the fossil record. New Phytologist, 207(2): 425–436.

Smith, S. A., Beaulieu, J. M., and Donoghue, M. J. 2010. An uncorrelated relaxed-clock analysis suggests an earlier origin for flowering plants. Proceedings of the National Academy of Sciences, 107(13): 5897–5902.

Stein, R. W., Brown, J. W., and Mooers, A. Ø. 2015. A molecular genetic time scale demonstrates cretaceous origins and multiple diversification rate shifts within the order Galliformes (Aves). Molecular Phylogenetics and Evolution, 92: 155–164.

Tank, D. C., Eastman, J. M., Pennell, M. W., Soltis, P. S., Soltis, D. E., Hinchliff, C. E., Brown, J. W., Sessa, E. B., and Harmon, L. J. 2015. Nested radiations and the pulse of angiosperm diversification: increased diversification rates often follow whole genome duplications. New Phytologist, 207(2): 454–467.

Thorne, J. L. and Kishino, H. 2002. Divergence time and evolutionary rate estimation with multilocus data. Systematic Biology, 51(5): 689–702.

Tong, K. J., Duchene, S., Ho, S. Y., and Lo, N. 2015. Comment on phylogenomics resolves the timing and pattern of insect evolution. Science, 349(6247): 487–487.

Warnock, R. C. M., Yang, Z., and Donoghue, P. C. J. 2012. Exploring uncertainty in the calibration of the molecular clock. Biology Letters, 8(1): 156–159.

Warnock, R. C. M., Parham, J. F., Joyce, W. G., Lyson, T. R., and Donoghue, P. C. J. 2015. Calibration uncertainty in molecular dating analyses: there is no substitute for the prior evaluation of time priors. Proceedings of the Royal Society of London B: Biological Sciences, 282(1798): 20141013.

Warnock, R. C. M., Yang, Z., and Donoghue, P. C. J. 2017. Testing the molecular clock using mechanistic models of fossil preservation and molecular evolution. Proceedings of the Royal Society of London B: Biological Sciences, 284(1857).

Wickham, H. 2009. ggplot2: Elegant Graphics for Data Analysis. Springer-Verlag New York.

Worobey, M., Han, G.-Z., and Rambaut, A. 2014. A synchronized global sweep of the internal genes of modern avian influenza virus. Nature, 508(7495): 254–257.

Yang, Z. 2007. PAML 4: phylogenetic analysis by maximum likelihood. Molecular Biology and Evolution, 24(8): 1586–1591.

Yang, Z. and Rannala, B. 2006. Bayesian estimation of species divergence times under a molecular clock using multiple fossil calibrations with soft bounds. Molecular Biology and Evolution, 23(1): 212–226.

Zeng, L., Zhang, Q., Sun, R., Kong, H., Zhang, N., and Ma, H. 2014. Resolution of deep angiosperm phylogeny using conserved nuclear genes and estimates of early divergence times. Nature Communications, 5: 4956.

Zhang, C., Stadler, T., Klopfstein, S., Heath, T. A., and Ronquist, F. 2015. Total-evidence dating under the fossilized birth-death process. Systematic Biology, 65(2): 228–249.

Zhu, T., Dos Reis, M., and Yang, Z. 2015. Characterization of the uncertainty of divergence time estimation under relaxed molecular clock models using multiple loci. Systematic Biology, 64(2): 267–280.

